## Supplemental File for "Zinc stimulation of phytoplankton in a low carbon dioxide, coastal Antarctic environment: evidence for the Zn hypothesis"

**Supplementary Table 1. Summary of zinc incubation experiments in marine environments.** Note that all experiments focus on phytoplankton populations with the exception of Fukada et al (2000) and Mazzotta et al. (2021) which focus on bacterial heterotrophy. This list represents our knowledge of the literature but may be incomplete.

| Study | Location | Zn Treatment Replication | $pCO_2$ Data | Dissolved Environmental Zn Data | Zn Response |
|---|---|---|---|---|---|
| Scharek et al. 1997[1] | Southern Ocean | Zn n=2, control n=1 | No | No | Negative/"very small" |
| Fukada et al., 2000[2] | North Pacific | Integrated water column leucine aminopeptidase activity (0-100m) | No | Yes | Leucine aminopeptidase activity positively correlated with dZn |
| Cochlan et al. 2002[3] | Western Ross Sea | None (n=1 +Zn, +Zn+Fe) | No | No | Increase in specific uptake of $NO_3^-$ |
| Coale et al. 2003[4] | Ross Sea, Southern Ocean | None (n=1 +Zn, +Zn+Fe) | No | Yes | Negative/"very small" |
| Crawford et al. 2003[5] | Northeast Pacific Ocean | Triplicate (n=3) | Yes | No | "Slightly but significantly" altered Chl, nitrate and phosphate |
| Franck et al. 2003[6] | Costa Rica Dome | Triplicate (n=3) | No | No | Secondary Zn limitation after Fe in diatom microscopy data |
| Ellwood 2004[7] | Subantarctic | Duplicates (n=2) | No | Yes | Negative |
| Cullen et al. 1999; Cullen and Sherrell 2005[8,9] | Coastal California | Duplicates (n=2) | No | No | Enhanced Cd:P in low $pCO_2$ treatments, decrease in Cd:P with Zn addition |
| Jakuba et al. 2012[10] | North Pacific | Singlicate with timepoints sacrificed | No | Yes | Primary, no additive effect with iron. |
| Dreux Chappell et al. 2016[11] | Costa Rica Dome | Triplicate (n=3) | No | Yes | Secondary limitation with Si |
| Sharma et al. 2020[12] | Eastern Arabian Sea | Triplicate (n=3) | Yes | No | Negative/ "insignificant effects" |
| Mazzotta et al. 2021[13] | Equatorial Pacific | Quintuplicate (n=5) sediment trap incubations | No | Yes in separate study (Cohen et al., 2021) | Enhancement of alkaline phosphatase activity with Zn addition in sediment trap samples |
| This study | Terra Nova Bay, Ross Sea | Triplicate (n=3) | Yes | Yes | Primary and Secondary limitation. Independent validation by Zn biomarkers. |

**Supplementary Table 2**. Station metadata for the NBP18-01 cruise. Stations at which total Zn uptake rates were determined are indicated by asterisks (*).

| Station | Latitude (°N) | Longitude (°E) | Sampling Date (yyyy-mm-dd hh:mm) |
|---|---|---|---|
| 4* | -72.751 | -116.001 | 2017-12-30 01:23 |
| 10 | -73.054 | -129.988 | 2018-01-03 03:39 |
| 11* | -74.047 | -133.764 | 2018-01-03 19:58 |
| 15* | -75.864 | -151.918 | 2018-01-05 14:51 |
| 20* | -76.714 | 179.819 | 2018-01-08 02:00 |
| 22* | -75.013 | 165.358 | 2018-01-09 15:56 |
| 25 | -75.293 | 163.914 | 2018-01-11 01:26 |
| 27* | -74.987 | 165.890 | 2018-01-11 16:05 |
| 29* | -76.001 | 172.997 | 2018-01-16 03:00 |
| 31 | -77.295 | 175.390 | 2018-01-17 04:39 |
| 32* | -76.750 | 172.000 | 2018-01-17 19:19 |
| 34 | -77.147 | 168.503 | 2018-01-23 22:59 |
| 35* | -76.231 | 168.769 | 2018-01-26 20:17 |
| 41* | -74.833 | 165.002 | 2018-01-29 00:41 |
| 46* | -74.742 | 165.287 | 2018-01-31 21:32 |
| 50 | -74.741 | 165.488 | 2018-02-02 21:17 |
| 52* | -75.000 | 164.005 | 2018-02-03 21:43 |
| 57* | -74.879 | 164.482 | 2018-02-06 20:12 |
| 60 | -74.959 | 164.739 | 2018-02-08 20:06 |
| 62* | -74.999 | 169.491 | 2018-02-09 19:27 |
| 67* | -76.454 | 167.919 | 2018-02-11 19:19 |
| 70 | -74.744 | 170.374 | 2018-02-13 21:25 |
| 72* | -74.800 | 164.395 | 2018-02-14 22:28 |
| 76* | -74.799 | 164.597 | 2018-02-16 20:15 |
| 78 | -74.696 | 164.796 | 2018-02-17 20:34 |
| 79* | -74.757 | 164.356 | 2018-02-18 19:29 |

**Supplementary Table 3**. Summary of ANOVA statistics for T6 incubation treatments comparing measured parameters. Significant differences among groups were found using one-way ANOVA and post-hoc Dunnett test (*** p < 0.001, ** p < 0.01, * p < 0.05, . p < 0.1). Chl *a*, chlorophyll *a;* $DIC_T$, total dissolved inorganic carbon; chl *b*, chlorophyll *b*; Prasino, prasinoxanthin; fuco, fucoxanthin; 19’hex, 19'-hexanoyloxyfucoxanthin; chl c3, chlorophyll c3. NA, no statistically significant difference. All parameters were measured in biological triplicate.

| Treatment (T6) | Chl *a* | $DIC_T$ | Chl *b* | Prasino | Fuco | 19’hex | Fuco:hex | Hex:Chl c3 | Bacterial abundance |
|---|---|---|---|---|---|---|---|---|---|
| +Fe vs Ctrl | *p* = 9.5e-5 (***) | *p* = 5.3e-6 (***) | *p* = 4.0e-3 (**) | *p* = 2.9e-2 (*) | NA | *p* = 8e-3 (**) | *p* = 4.2e-4 (***) | *p* = 2.0e-4 (***) | *p* = 9.1e-4 (***) |
| +Zn vs Ctrl | *p* = 1.1e-2 (*) | *p* = 5.0e-6 (***) | *p* = 8.0e-2 (.) | *p* = 7.4e-3 (**) | NA | NA | NA | *p* = 0.02520 (*) | NA |
| +FeZn vs Ctrl | *p* = 1.3e-7 (***) | *p* = 2.2e-16 (***) | *p* = 4.0e-4 (***) | *p* = 5.7e-2 (.) | NA | NA | *p* = 2.7e-3 (**) | *p* = 1.9e-4 (***) | *p* = 6.3e-4 (***) |
| +FeZn vs +Fe | *p* = 3.4e-2 (*) | *p* = 4.4e-3 (**) | NA | NA | NA | *p* = 6.0e-2 (.) | NA | NA | NA |

**Supplementary Table 4.** Representative proteins of interest, reference organism and IDs.

| Protein of interest | Reference organism | Protein ID |
|---|---|---|
| ZCRP-A | *Thalassiosira pseudonana* CCMP1335 | 3054 (JGI Thaps3[*]) |
| ZCRP-B | *Thalassiosira pseudonana* CCMP1335 | 938 (JGI Thaps3_bd[**]) |
| RUBISCO | *Phaeodactylum tricornutum* CCMP632 | AAF07200.1 (NCBI) |
| ISIP1A | *Thalassiosira oceanica* CCMP1005 | K0RCT3 (Uniprot) |
| ISIP2A | *Phaeodactylum tricornutum* CCMP632 | B7FYL2 (Uniprot) |
| ISIP3 | *Phaeodactylum tricornutum* CCMP632 | B7G4H8 (Uniprot) |
| ZIP | *Phaeodactylum tricornutum* CCMP632 | 46780 (JGI Phatr2[†]) |
| CDCA | *Thalassiosira pseudonana* CCMP1335 | 25840 (JGI Thaps3[*]) |

[*]Joint Genome Institute (JGI) Thaps3 database (https://mycocosm.jgi.doe.gov/Thaps3/Thaps3.home.html)

[**]Joint Genome Institute Thaps3_bd database (https://mycocosm.jgi.doe.gov/Thaps3_bd/Thaps3_bd.home.html

[†]Joint Genome Institute CCAP 1055/1 v2.0 Phatr2, all models database (https://mycocosm.jgi.doe.gov/Phatr2/Phatr2.home.html)

**Supplementary Table 5**. Particulate Zn:P ratios measured in *T. pseudonana* cultured at various concentrations of Zn (log [Zn′]) (Sunda and Huntsman 2005) compared to particulate Zn:P ratios measured at Station 27 of research cruise NBP18-01. Zn:C measurements by Sunda and Huntsman 2005 were converted to Zn:P using the Redfield ratio. Data is plotted in Supplementary Figure 6. Zn′ refers to the sum of inorganic Zn metal complexes.

[+]Particulate P data measured directly in this study

*Particulate P from Sunda 2005 data was estimated by converting particulate C measurements to P using the Redfield ratio (106C:1P).

| **Particulate Zn:P ratios at Station 27** | | | | | | | |
|---|---|---|---|---|---|---|---|
| Depth (m) | | | Particulate Zn (mol) | | Particulate P (mol)[+] | Zn:P (mol:mol) | |
| 100 | | | 1E-11 | | 8E-08 | 1E-04 | |
| 50 | | | 1E-11 | | 1E-07 | 1E-04 | |
| 25 | | | 3E-11 | | 2E-07 | 2E-04 | |
| 10 | | | 4E-11 | | 2E-07 | 2E-04 | |
| **Cellular Zn:P ratios of cultured *T. pseudonana* from Sunda and Huntsman 2005, Table 1.** | | | | | | | |
| Experiment | Log [Zn’] | pH | Particulate Zn (mol) | Particulate C (mol) | Particulate P (mol)* | Zn:P (mol:mol) | Growth rate (d$^{-1}$) |
| 1 | -12.05 | 8.2 | 3.3E-05 | 22 | 0.21 | **2E-04** | **0.1** |
| 1 | -11.47 | 8.2 | 5.2E-05 | 15 | 0.14 | 4E-04 | 1.23 |
| 1 | -10.87 | 8.2 | 1.2E-04 | 15 | 0.14 | 9E-04 | 1.62 |
| 1 | -10.36 | 8.2 | 1.5E-04 | 15 | 0.14 | 1E-03 | 1.76 |
| 1 | -9.82 | 8.2 | 1.8E-04 | 15 | 0.14 | 1E-03 | 1.76 |
| 1 | -11.96 | 9 | 3.8E-05 | 15 | 0.14 | 3E-04 | 0.51 |
| 1 | -11.38 | 9 | 8.5E-05 | 15 | 0.14 | 6E-04 | 1.12 |
| 1 | -10.78 | 9 | 2.0E-04 | 15 | 0.14 | 1E-03 | 1.28 |
| 1 | -10.27 | 9 | 4.1E-04 | 15 | 0.14 | 3E-03 | 1.42 |
| 1 | -9.73 | 9 | 5.0E-04 | 15 | 0.14 | 4E-03 | 1.49 |
| 2 | -11.82 | 8.2 | 4.9E-05 | 15 | 0.14 | 3E-04 | 0.64 |
| 2 | -11.82 | 8.2 | 4.0E-05 | 15 | 0.14 | 3E-04 | 0.79 |
| 2 | -10.87 | 8.2 | 1.3E-04 | 15 | 0.14 | 9E-04 | 1.45 |
| 2 | -9.82 | 8.2 | 2.0E-04 | 15 | 0.14 | 1E-03 | 1.51 |
| 2 | -11.71 | 9 | 5.8E-05 | 15 | 0.14 | 4E-04 | 0.73 |
| 2 | -11.71 | 9 | 5.8E-05 | 15 | 0.14 | 4E-04 | 0.78 |
| 2 | -10.76 | 9 | 1.9E-04 | 15 | 0.14 | 1E-03 | 1.39 |
| 2 | -10.76 | 9 | 2.1E-04 | 15 | 0.14 | 1E-03 | 1.38 |
| 2 | -9.71 | 9 | 4.3E-04 | 15 | 0.14 | 3E-03 | 1.44 |
| 2 | -9.71 | 9 | 5.0E-04 | 15 | 0.14 | 4E-03 | 1.42 |
| 4 | -11.82 | 8.2 | 2.8E-05 | 18.1 | 0.21 | 3E-04 | 0.72 |
| 4 | -11.32 | 8.2 | 5.5E-05 | 15.2 | 0.13 | 4E-04 | 1.45 |
| 4 | -10.82 | 8.2 | 1.3E-04 | 14.7 | 0.14 | 9E-04 | 2 |
| 4 | -10.32 | 8.2 | 2.2E-04 | 15.1 | 0.14 | 2E-03 | 2.03 |
| 4 | -9.82 | 8.2 | 2.6E-04 | 15.6 | 0.14 | 2E-03 | 2.04 |

**Supplementary Table 6**. Reference seawater comparisons using the 2009 GEOTRACES coastal surface seawater (GSC) standard.

| Metal | This study (n = 8) (nM) | GEOTRACES GSC consensus (nM) |
|---|---|---|
| Fe | 1.6 ± 0.23 | 1.6 ± 0.12 |
| Zn | 1.4 ± 0.23 | 1.5 ± 0.10 |
| Cd | 0.4 ± 0.01 | 0.4 ± 0.02 |
| Cu | 1.3 ± 0.05 | 1.1 ± 0.15 |
| Ni | 4.2 ± 0.07 | 4.5 ± 0.21 |
| Mn | 2.1 ± 0.37 | 2.2 ± 0.08 |

### Supplementary Figures

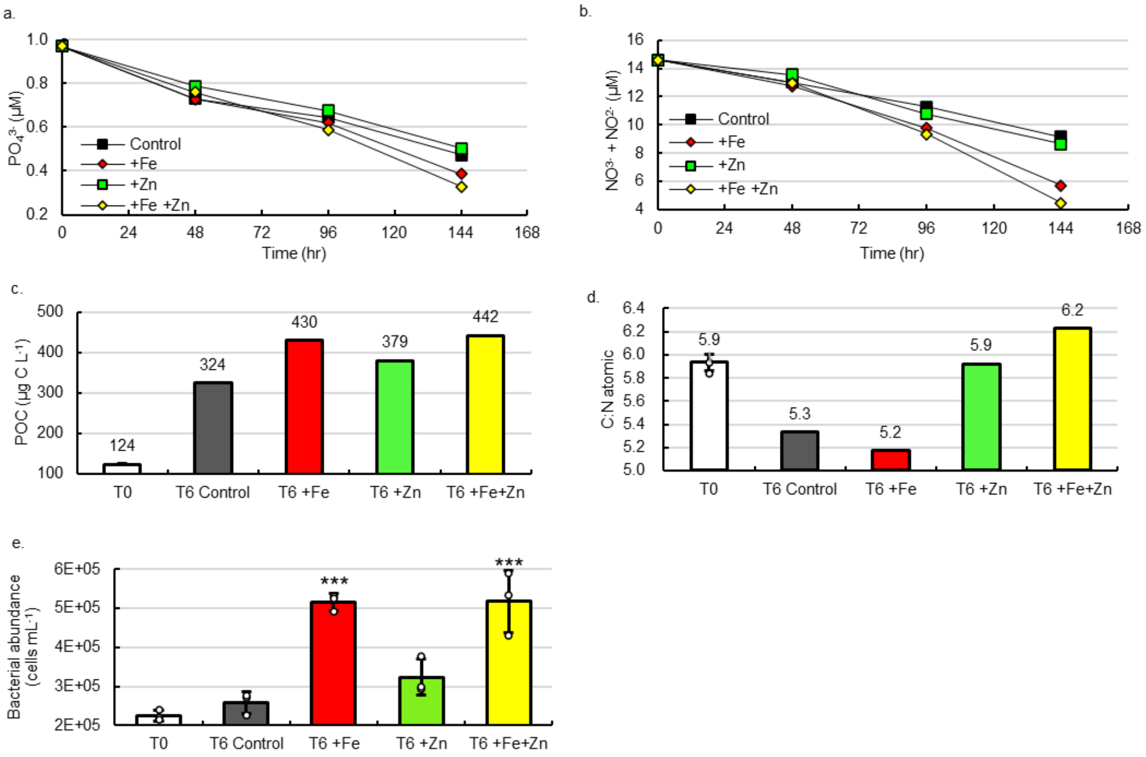


**Supplementary Figure 1. Additional parameters measured in shipboard bottle incubations**. Time course decreases in (**a**) phosphate and (**b**) nitrite + nitrate. (**c**) total POC of biomass in each treatment at T6. (**d**) the atomic carbon:nitrogen (C:N) ratio of biomass in each treatment at T6, and (**e**) bacterial abundance. Significant differences among groups were found using one-way ANOVA and post-hoc Dunnett test (*** $p < 0.001$, ** $p < 0.01$, * $p < 0.05$). Error bars are the standard deviation of biological triplicates (n=3) with individual data points overlaid (white circles). Macronutrients, POC, and PON were measured in singlicate (n=1) from pooled biological triplicates.

a.

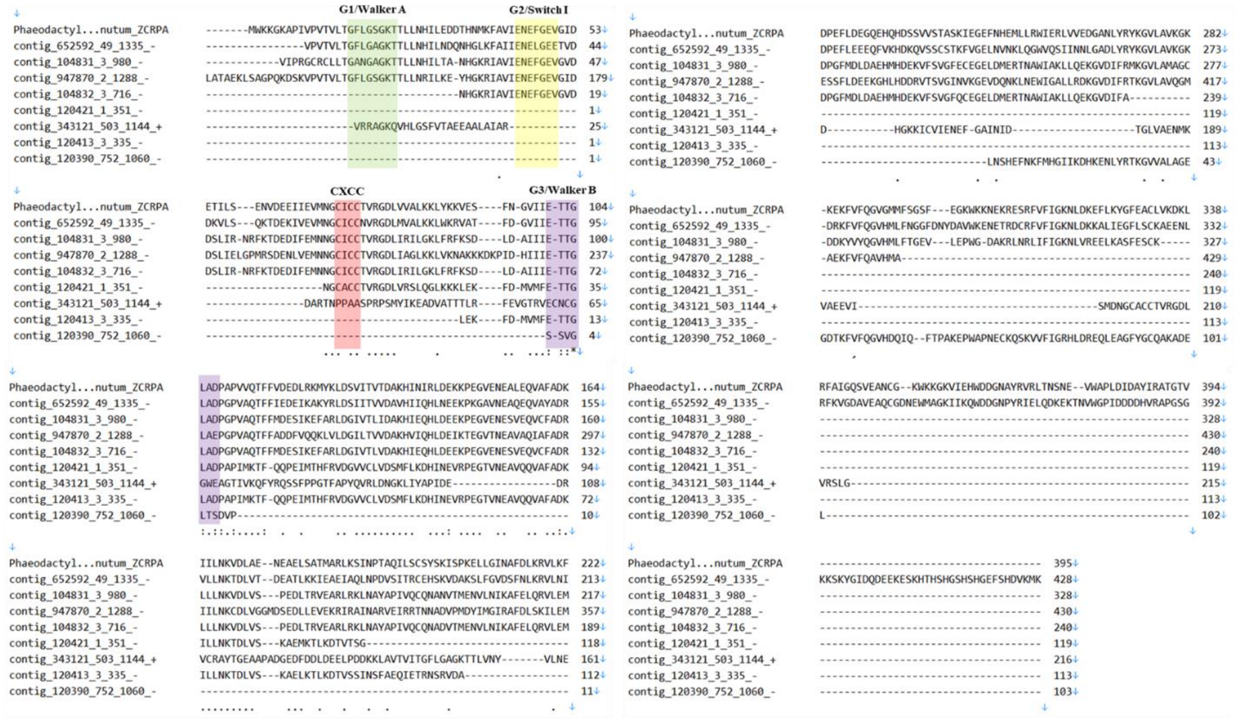


b.

| orf_id | Hit | E value | % ID |
|---|---|---|---|
| contig_652592_49_1335_- | Phaeodactylum_tricornutum_ZCRPA | 4.75E-162 | 56.266 |
| contig_104831_3_980_- | Phaeodactylum_tricornutum_ZCRPA | 6.51E-110 | 50.311 |
| contig_947870_2_1288_- | Phaeodactylum_tricornutum_ZCRPA | 5.00E-86 | 49.488 |
| contig_104832_3_716_- | Phaeodactylum_tricornutum_ZCRPA | 5.25E-80 | 48.954 |
| contig_120421_1_351_- | Phaeodactylum_tricornutum_ZCRPA | 1.62E-32 | 51.818 |
| contig_343121_503_1144_+ | Phaeodactylum_tricornutum_ZCRPA | 3.24E-23 | 60.526 |
| contig_120413_3_335_- | Phaeodactylum_tricornutum_ZCRPA | 5.54E-22 | 41.964 |
| contig_120390_752_1060_- | Phaeodactylum_tricornutum_ZCRPA | 7.76E-13 | 41.111 |

**Supplementary Figure 2. Sequence alignments of ZCRP-A peptides detected in T6 incubation biomass. (a)** Sequence alignment of the *Phaeodactylum tricornutum* ZCRP-A protein compared to all ZCRP-A proteins detected in T6 incubation biomass. Alignment was generated using the MUSCLE algorithm with default parameters within MEGA11. Four conserved GTPase (G1/Walker A, G2/SwitchI, CXCC metal binding, and G3/Walker B) are labeled. (**b**) E values and % identities of the identified proteins with significant sequence similarity to *P. tricornutum* ZCRP-A aligned above.

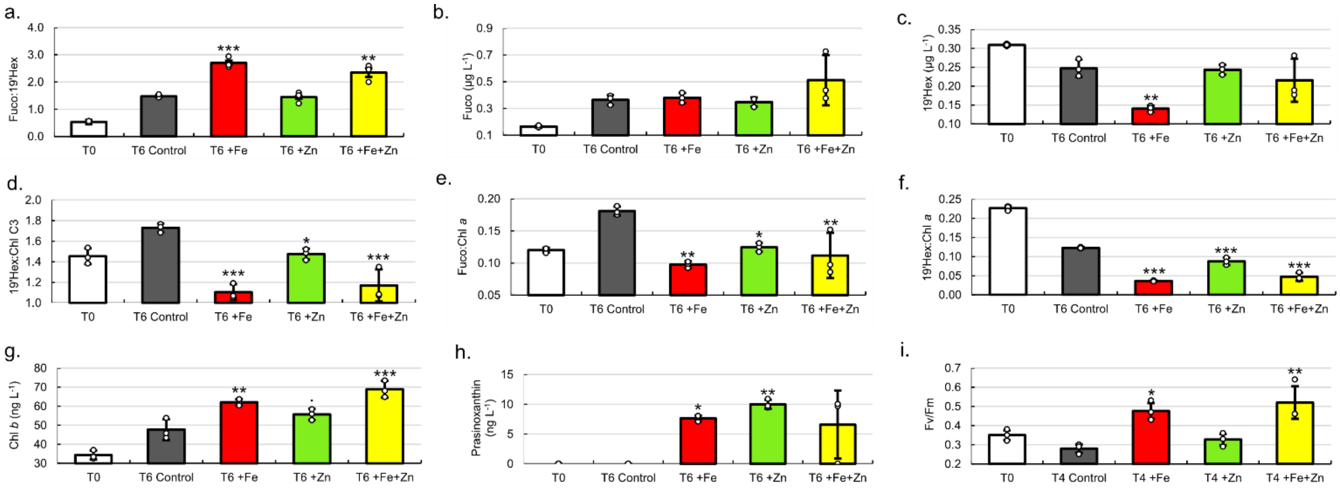


**Supplementary Figure 3. Pigment analysis of T6 incubations**. (**a**) ratio of fucoxanthin: 19'-Hex, (**b**) fucoxanthin, (**c**) 19-hexanoyloxyfucoxanthin (19'-Hex), (**d**) ratio of 19'-Hex: chl c3, (**e**) ratio of fucoxanthin: chlorophyll *a*, (**f**) ratio of 19'-Hex: chlorophyll *a*, (**g**) chlorophyll *b*, (**h**) prasinoxanthin, and (**i**) maximum quantum efficiency (Fv/Fm) among treatments at T4. Significant differences among groups were found using one-way ANOVA and post-hoc Dunnett test (*** $p < 0.001$, ** $p < 0.01$, * $p < 0.05$, . $p < 0.1$). Data with error bars are presented as mean values ± the standard deviation of biological triplicates (n=3) with individual data points overlaid (white circles).

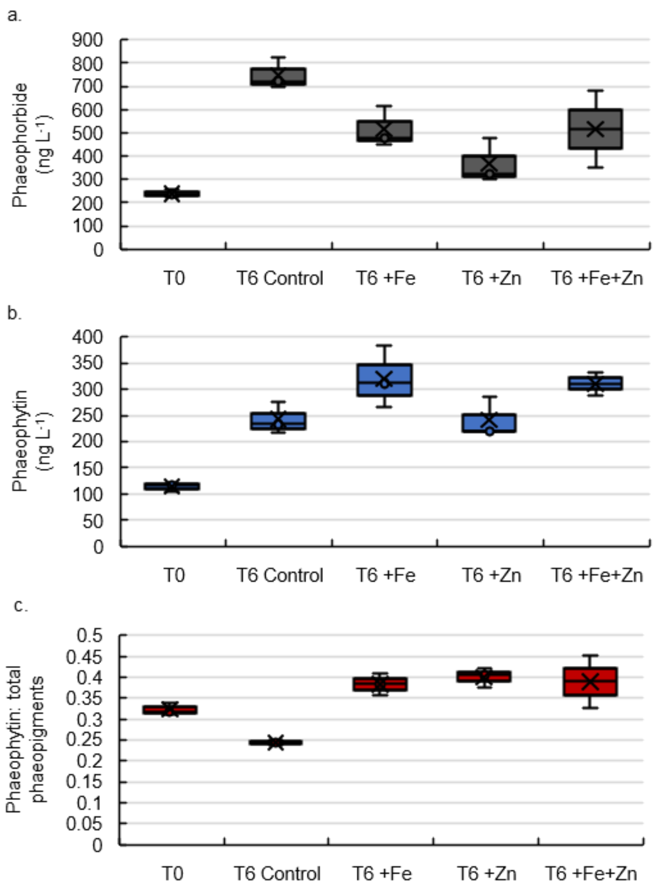


**Supplementary Figure 4.** Phaeopigments measured in T6 incubation biomass. Abundances of (**a**) phaeophorbide and (**b**) phaeophytin, and (**c**) the ratio of phaeophytin: total phaeopigments (the sum of phaeophorbide and phaeophytin). Error bars are the standard deviation of biological triplicates (n=3).

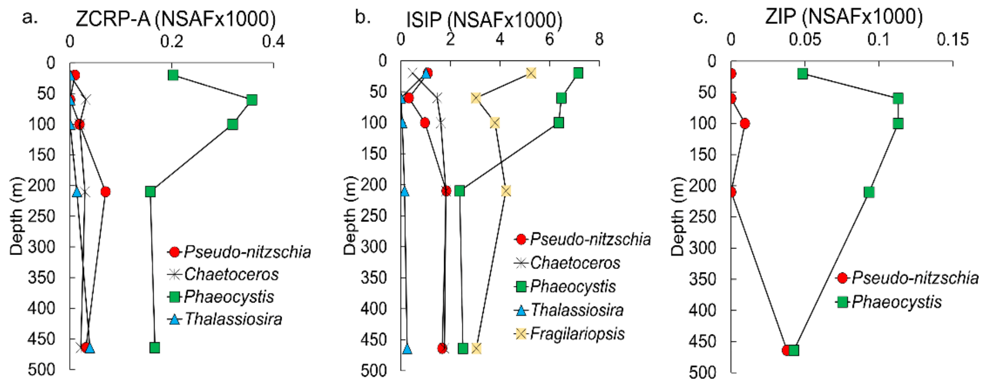


**Supplementary Figure 5. Depth profiles for proteins of interest at station 27 categorized by genus.** Depth profiles of NSAF-normalized protein spectral counts of (**a**) ZCRP-A, (**b**), iron starvation induced proteins (ISIPs), and (**c**) ZIPs summed by genus. Proteins assigned to an individual genus were summed across all size fractions (0.2, 3 and 51 μm). ISIPs are the combined spectral counts of ISIP1A, ISIP2A and ISIP3.

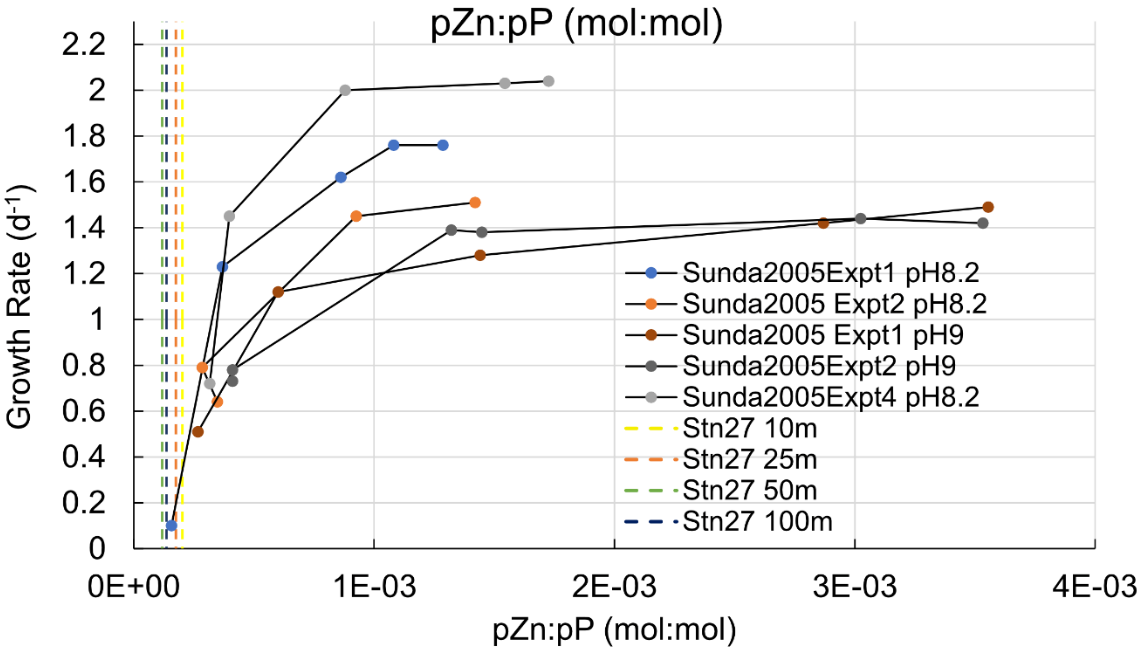


**Supplementary Figure 6. Comparison of particulate Zn: particulate P ratios measured in the water column at the experimental site to those ratios measured in culture studies of a Zn-limited diatom.** Ratio of particulate Zn (pZn) and particulate phosphorus (pP) (pZn:pP) measured in the upper water column at the study site (station 27; vertical dashed lines) compared to pZn:pP measurements in Zn limitation studies of the diatom *Thalassiosira pseudonana* in culture by Sunda and Huntsman 2005.

**Supplementary Information References**
